# Spatial conservation planning of forest genetic resources in a Mediterranean multi-refugial area

**DOI:** 10.1101/2023.11.06.565750

**Authors:** Elia Vajana, Marco Andrello, Camilla Avanzi, Francesca Bagnoli, Giovanni G. Vendramin, Andrea Piotti

## Abstract

The Kunming-Montreal Global Biodiversity Framework recognised the urgency of taking action to conserve intraspecific genetic diversity (IGD) as an insurance against habitat degradation and environmental change. Recent work suggests that 90-99% of IGD should be conserved to safeguard viability of future generations.

Here, we addressed such a conservation issue in three forest tree species in Italy: silver fir (*Abies alba* Mill.), Heldreich’s pine (*Pinus heldreichii* H. Christ), and pedunculate oak (*Quercus robur* L.). We used microsatellite markers to measure IGD of 36 (*A. alba*), 15 (*P. heldreichii*) and 25 (*Q. robur*) natural sites, including several putative glacial refugia. We developed a spatial conservation planning (SCP) analysis to quantify the genetic irreplaceability of each site and identify the minimum set coverage ensuring IGD protection. Finally, we compared SCP results with the contributions to allelic diversity within and between sites, total allelic diversity and private allelic richness.

We found that between 44% and 73% of sites were required to conserve 90-99% of the alleles, and that this conservation effort held even when targeting lower percentages of alleles to protect (50-75%). Glacial refugia were often included in the minimum set coverage, confirming theoretical and biogeographical expectations. Finally, sites with high genetic irreplaceability were found to have higher private allelic richness on average. These results are discussed in the light of the biogeographic history of the species studied and the current policies for the conservation of forest genetic resources.

## 1. Introduction

Intraspecific genetic diversity (IGD), encompassing genetic variation within and between populations, is a key component of ecosystem functioning as it underlies the species’ capacity to adapt to the environment, survive environmental change and provide human society with ecosystem services (Robuchon et al., 2022). Particularly, IGD of forest trees (*aka* forest genetic resources) contributes to climate regulation, water filtration, nutrient cycling, wood production, and coastal and soil protection against erosion (Alberto et al., 2013; Gougherty et al., 2021; Panagos et al., 2015; Schweitzer et al., 2011). The preservation of forest genetic resources lies at the core of sustainable forest management and conservation (Fady et al., 2016). Yet, human activities inducing global warming, habitat fragmentation and overharvesting are accelerating biodiversity decline worldwide, with massive losses in genetic diversity and unpredictable impacts on the biosphere (Exposito-Alonso et al., 2022; Hoban et al., 2023). Consequently, between 19 and 66% of the whole amount of IGD on Earth is estimated to go lost in the next decades unless specific actions are undertaken to halt genetic erosion and sustain evolutionary rescue (Hoban et al., 2023, 2021).

Among the long-term goals for 2050, the Kunming-Montreal global biodiversity framework formally recognized the need of conserving IGD and the urgency of putting in place actions to protect ‘the genetic diversity within and between populations of native, wild, and domesticated species to maintain their adaptive potential’ (Convention on Biological Diversity, 2022; Hoban et al., 2023). To this aim, conservation of 90-99% of extant IGD is usually advocated together with guaranteeing an effective population size >500 so as to reduce genetic erosion and prevent ‘future losses for all populations […] regardless of past losses’ (Frankham, 2022; Hoban et al., 2023).

Area-based conservation is among the approaches proposed to contrast genetic erosion (Hoban et al., 2023; Robuchon et al., 2021). However, budget constraints usually require choices to be made when attempting to preserve IGD with such a strategy (Marris, 2007; Vane-Wright et al., 1991; Weitzman, 1998). Then, scientifically based criteria were developed for setting spatial priorities for conservation, e.g., whether to prioritise genetically distinct vs. highly diverse populations (Fernandez-Fournier et al., 2021; Petit et al., 1998; Weitzman, 1993) or sites with endemic lineages rather than centres of radiation (Erwin, 1991; Robuchon et al., 2021). More recently, spatial conservation planning (SCP) was applied to optimise the selection of new conservation areas based on genetic information (Andrello et al., 2022; Nielsen et al., 2023a). In this case, spatially optimised solutions are found across the study area to minimise the costs required to achieve specific conservation objectives for IGD, a formulation of the SCP problem known as ‘minimum set coverage’ (Moilanen et al., 2009).

Here, we used SCP to identify areas that could maximise the protection of IGD in three keystone forest tree species characterised by a highly fragmented distribution in Italy – two mountainous conifers, i.e., silver fir (*Abies alba* Mil.) and Heldreich’s pine (*Pinus heldreichii* H. Christ), and a broadleaved lowland tree, the pedunculate oak (*Quercus robur* L.). Palaeobotanical and genetic evidence proved that the Mediterranean region provided temperate trees with climatically favourable refugial areas during the glacial periods characterising the Quaternary (Petit et al., 2003). Most glacial refugia were located in the Iberian, Italian and Balkan peninsulas, with populations inhabiting these areas generally showing high levels of genetic distinctiveness (Petit et al., 2003). This genetic feature makes forest tree populations from former glacial refugia candidate sites for conserving forest genetic resources, as they are likely to contain unique variation that may prove necessary to foster adaptive potential in the face of climate change (Petit et al., 2005). Here, we focused our analysis on a set of sites within a complex of refugial areas (Brewer et al., 2002; Petit et al., 2002; Piotti et al., 2017), where fragmentation is likely pushing peripheral populations towards an extinction vortex in concert with climate change (Bucci et al., 1997; Gibbs, 1997; Maiorano et al., 2013; Ruosch et al., 2016; Thomas et al., 2002).

The goals of the present work were (i) to identify priority sites for conserving IGD in silver fir, Heldreich’s pine and pedunculate oak in a multi-refugial area, and (ii) to investigate the consensus between priority sites identified through SCP and those found through other genetic parameters, namely allelic diversity and its decomposition into within-site and between-site contributions (Petit e al., 1998), and effective population size, which is related to genetic load and adaptive potential and has been formally adopted by the Convention on Biological Diversity (2022) to monitor intraspecific genetic diversity.

## 2. Materials and methods

### 2.1. Sampling and genotyping

We relied on genetic data from 36 natural populations of *A. alba* (seven of which were specifically characterised for this study, resulting in 1769 trees in total, 49 on average per site; **Supplementary Table 1**), 15 natural populations of *P. heldreichii* (515 trees, 34 on average), and 25 natural populations of *Q. robur* (745 trees, 30 on average) to feed SCP analyses (**Figure 1**). The large representativeness of IGD was guaranteed by densely covering the local distribution of the species, and by including sampling sites from all known putative glacial refugial areas.

**Table 1.** Median number of sites needed to conserve 50%, 75%, 90%, 99%, and 100% of IGD across 100 replicates in *Abies alba, Pinus heldreichii* and *Quercus robur*.

|  | Protection levels |  |  |  |  |
| --- | --- | --- | --- | --- | --- |
|  | 50% | 75% | 90% | 99% | 100% |
| ***A. alba* (36)** | 14 ($\pm 1$)<sup>a</sup> | 15 (ND)<sup>b</sup> | 16 (ND) | 16 (ND) | 16<sup>c</sup> |
| ***P. heldreichii* (15)** | 8 ($\pm 1$) | 10 ($\pm 1$) | 11 (ND) | 11 (ND) | 11 |
| ***Q. robur* (25)** | 15 ($\pm 1$) | 17 ($\pm 1$) | 18 (ND) | 18 (ND) | 18 |

<sup>a</sup> Median absolute deviation (MAD), i.e., an estimate of the variability around the number of sites in the replicates. Each MAD was calculated as the median of the absolute deviations from the number of sites. <sup>b</sup> No deviation from the number of sites observed. <sup>c</sup> No variation was associated with the 100% level as a unique analysis was performed in this scenario. The total number of sampled sites is given in brackets next to each species name (first column).

**Figure 1.**
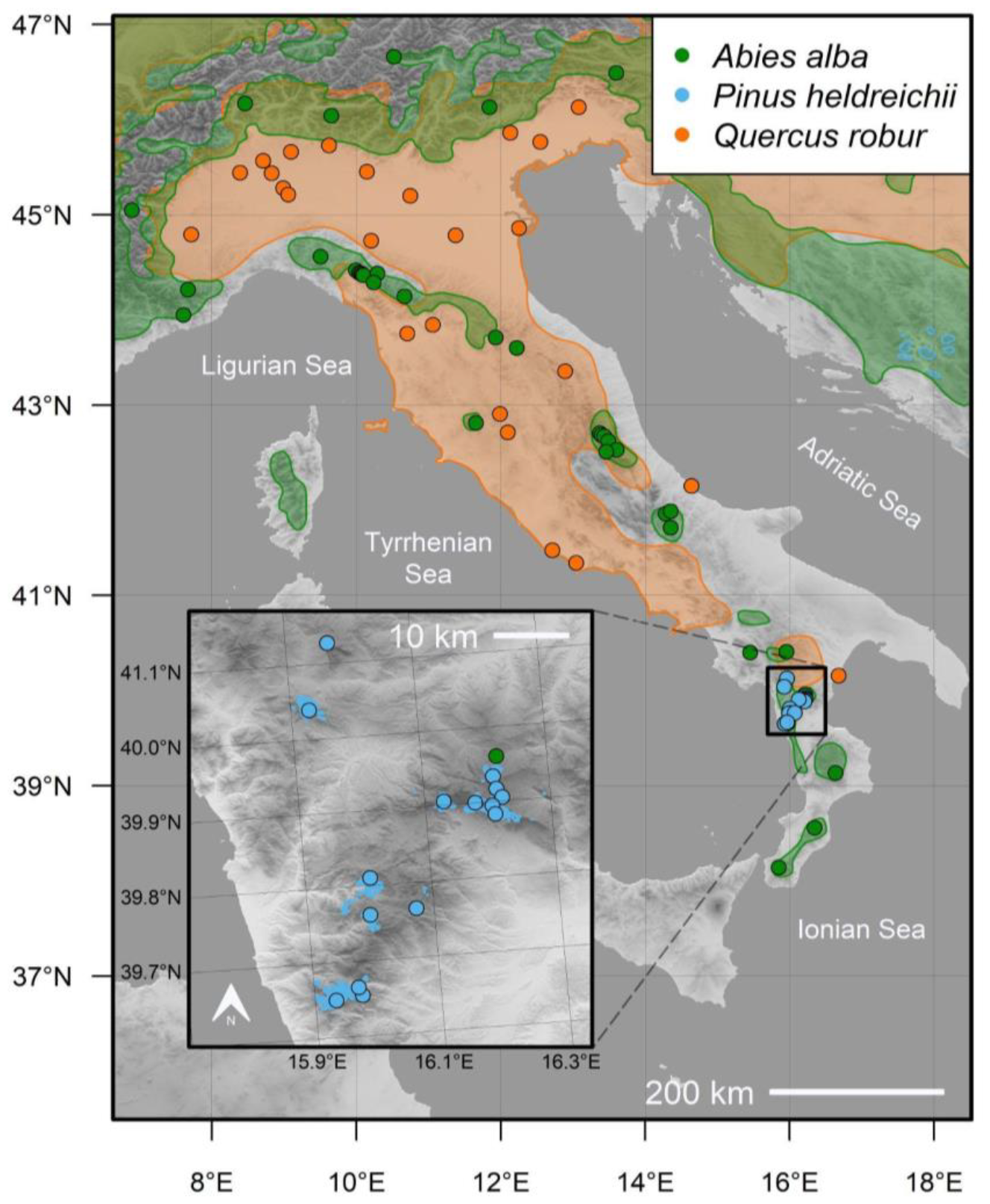
Sampling sites: *Abies alba* (green points), *Pinus heldreichii* (blue points), and *Quercus robur* (orange points). Approximate species distributions (Caudullo et al., 2017) are also shown in the background with the same colour scheme used to indicate sampling sites (note: *A. alba* and *Q. robur* distributions are rather imprecise representations of the actual distribution of these species in the study area). The inset map zooms over the Pollino National Park, which encircles the distribution of Heldreich’s pine in Italy. Shades of grey indicate elevation.

*Abies alba* was genotyped at 16 nuclear short sequence repeats (nSSRs), resulting in 145 alleles (Piotti et al., 2017; Santini et al., 2018), *P. heldreichii* at 12 nSSRs (89 alleles; Piovesan et al., 2023), and *Q. robur* at 16 nSSRs (245 alleles; Avanzi et al., 2023). The details about genotyping and standard population genetic analyses can be found in the above-mentioned references.

### 2.2. Spatial conservation planning

The SCP analysis was run separately for each species, as there were no common sampling sites to help identify priority sites shared by more than one species. In each species, each allele scored was considered as a distinct conservation feature (see the ‘AL’ method in Andrello et al., 2022). The spatial distribution of each allele was coded as presence/absence in each site. The number of selected sites was considered as a simple proxy for total conservation cost (Paz-Vinas et al., 2018; von Takach et al., 2021). The objective was to find solutions for conserving low (50%), moderate (75%) and high (90%, 99% and 100%) proportions of protected alleles. These proportions of protected alleles were interpreted as levels of protected IGD (Paz-Vinas et al., 2018), and included the 90% objective proposed in the first draft of the post-2020 GBF (CBD, 2021), as well as the 99% objective proposed more recently (Frankham, 2022; Hoban et al., 2023).

The SCP was run under the ‘minimum set coverage’ formulation to minimise the total conservation cost needed to achieve a given level of IGD protection (Andrello et al., 2022). This was operationalized as finding the minimum set of sites ensuring that the given proportion of alleles (i.e., 50%, 75%, 90%, 99% or 100%) was adequately protected. An allele was considered as adequately protected if at least 30% of its occurrence sites were included in the solution as an insurance against catastrophic events (Lefèvre et al., 2020; Wilson et al., 2009). For the 50%, 75%, 90% and 99% IGD levels, we performed 100 independent replicates, each based on a random selection of alleles to be protected. We calculated the genetic irreplaceability (GI) of each site as the number of times each site was selected across the 100 replicates (Pressey et al., 1994). For the 100% IGD level, a single analysis was performed as all the alleles were included by definition, which allowed the prioritisation problem to find a unique, optimal set of sites needed to conserve all the alleles. All SCP analyses were performed using the R package ‘prioritizer’ v. 7.2.2 (Hanson et al., 2022) and solved within 10% optimality using the ‘Gurobi’ solver v. 9.5.2 (Gurobi Optimization, LLC, 2023).

### 2.3. Relationship between genetic irreplaceability and genetic diversity parameters

We studied the relationship between GI and five genetic parameters commonly used to prioritise populations for conservation: the relative site-specific contributions to (1) within-site (*A*_S_) and (2) between-site (*D*_A_) allelic diversity (the mean number of alleles occurring in the sites under study, and the mean number of alleles occurring in a site and absent in another, respectively), and (3) total allelic diversity (*A*_T_; the sum of *A*_S_ and *D*_A_), as well as (4) private allelic richness (*PAr*; the number of unique alleles in a given site) and (5) effective population size (*Ne*; the size of a Wright-Fisher’s idealised population displaying the same rate of inbreeding or drift observed in the population under study, roughly corresponding to the number of reproductive individuals contributing to next generations; Jamieson and Allendorf, 2012). Sites mostly contributing to *A*_T_ can be identified by their relative contribution to *A*_S_ and *D*_A_ (Petit et al., 1998). *A*_T_, *A*_S_ and *D*_A_ were calculated for all sites in each species with the ‘metapop2’ software (López‐Cortegano et al., 2019). *PAr* was estimated with ‘HP-RARE’ (Kalinowski, 2005). Rarefaction was used to account for uneven sample sizes in the calculation of *A*_T_, *A*_S_, *D*_A_, and *PAr* (El Mousadik and Petit, 1996). *Ne* was estimated using the linkage disequilibrium method (Waples and Do, 2010) implemented in ‘NeEstimator’ v. 2.1 (Do et al., 2014) and screening out alleles with a frequency <2%.

The relationship between GI and genetic parameters was assessed based on the GI values obtained when protecting 99% of alleles. Considering the distribution of GI values, sites were classified to be at low/high GI, using a threshold of ≤20% (low GI) and ≥80% (high GI), respectively. Associations between levels of GI and genetic parameters were tested through either the Welch or Student’s two sample *t*-test, depending on whether the variances of the groups resulted to be unequal or equal based on the Levene’s test. To reduce inflation in the familywise type I error due to multiple testing, the Bonferroni correction was applied to the *p*-values of the tests involving the same species. A test was deemed to be statistically significant if its adjusted *p*-value was equal to or lower than the nominal significance threshold (α) of 0.05. The lower bound of the confidence intervals around the *Ne* estimates was used in the tests to avoid dealing with infinite *Ne* values. All operations were performed in the R programming environment v. 4.3.2 (R Core Team, 2023).

## 3. Results

### 3.1. Spatial conservation planning

In all species, the number of sites needed for conserving 100% IGD was less than the total number of sampled sites, and was equal to 16 (44% of sampled sites) in *A. alba*, 11 (73%) in *P. heldreichii*, and 18 (72%) in *Q. robur*. For conserving lower levels of IGD, the median number of sites needed was equal or slightly lower with almost no variation among the replicates (**Table 1**).

Regardless of the number of alleles to be protected in the SCP analysis, we found that ∼98% of the replicates conserved at least 90% of all alleles present in the dataset (**Figure 2**). All replicates protected >90% of alleles present in the dataset when including 75% of alleles in the SCP analysis. Overall, the SCP analyses systematically ensured protection for higher numbers of alleles than those included as conservation features.

**Figure 2.**
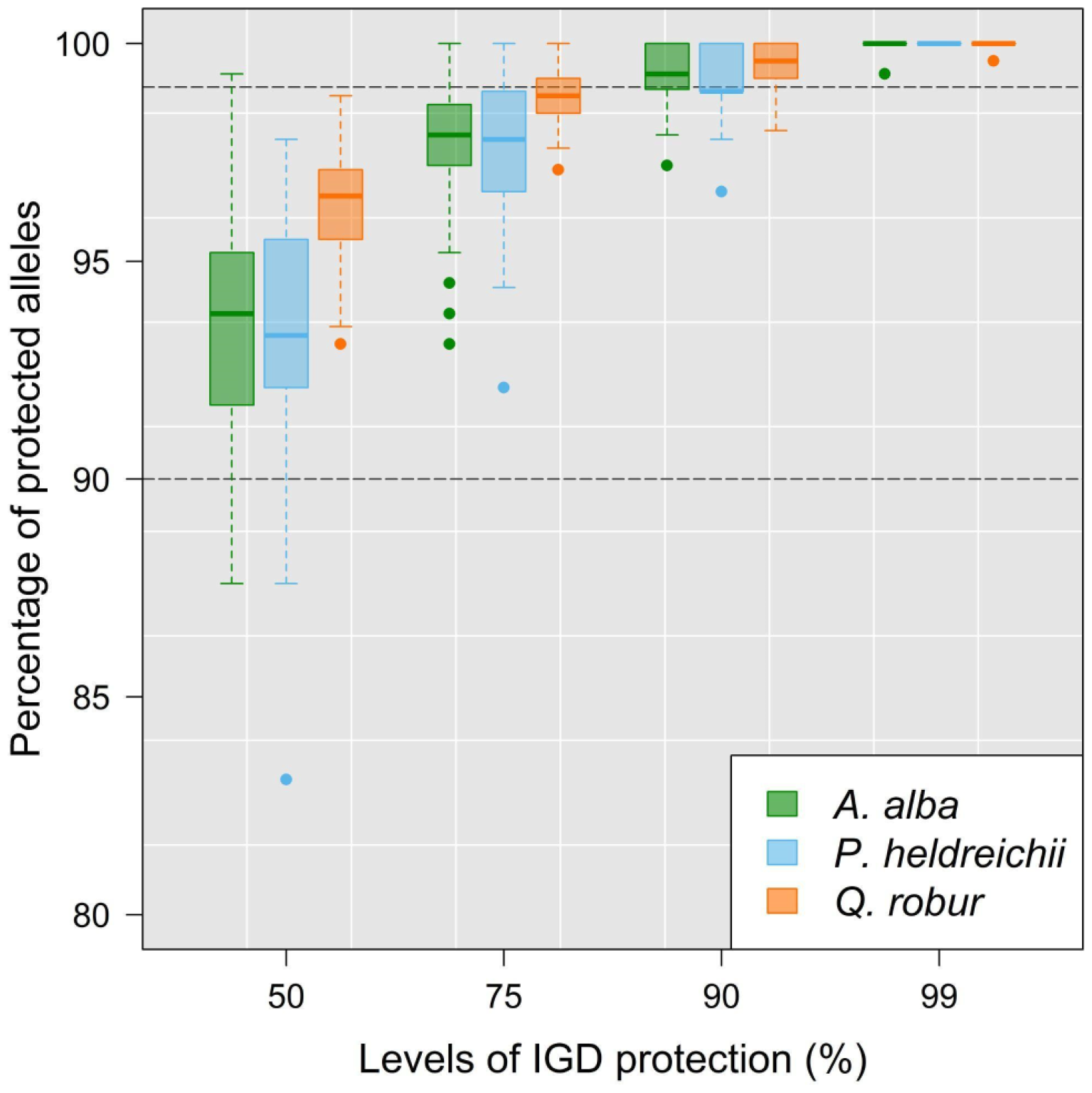
Variation in the percentage of alleles protected in at least 30% of their sites of occurrence as a function of levels of IGD secured (50%, 75%, 90%,99% and 100%). The proportion of alleles adequately protected is represented separately for each species.

For *A. alba*, a bimodal spatial pattern of GI appeared and four sites, two in the northern Apennines and two in the southern Apennines, were irreplaceable (i.e., GI=100%) when 90% of the alleles were conserved (**Figure 3a**). When targeting the 99% level, 11 sites became irreplaceable and were more evenly distributed across the Italian peninsula (**Figure 3b**). For *P. heldreichii*, 11 sites showed GI≥88% when conserving ≥90% of alleles, including one irreplaceable site on the southern margins (**Figure 3c**). When 99% of the alleles were conserved, all such 11 sites became irreplaceable (**Figure 3d**). Irreplaceable sites were located at short distances (<10 Km apart), and included a group of five close, geographically central sites, and three sites on the southern margins. For *Q. robur*, we found three irreplaceable sites even when conserving 75% of alleles (result not shown), including the southernmost site in Italy (‘Bosco Pantano’). When protecting 90% of alleles (**Figure 3e**), four sites were found to be irreplaceable, including a new site in the Po valley (northern Italy). In this case, we found 18 sites with GI≥68%, and seven sites with GI≤27%. The same sites at higher GI were confirmed when conserving 99% of the alleles, when we observed a GI higher than 96% and lower than 5% in sites at high vs. low GI, respectively (**Figure 3f**).

**Figure 3.**
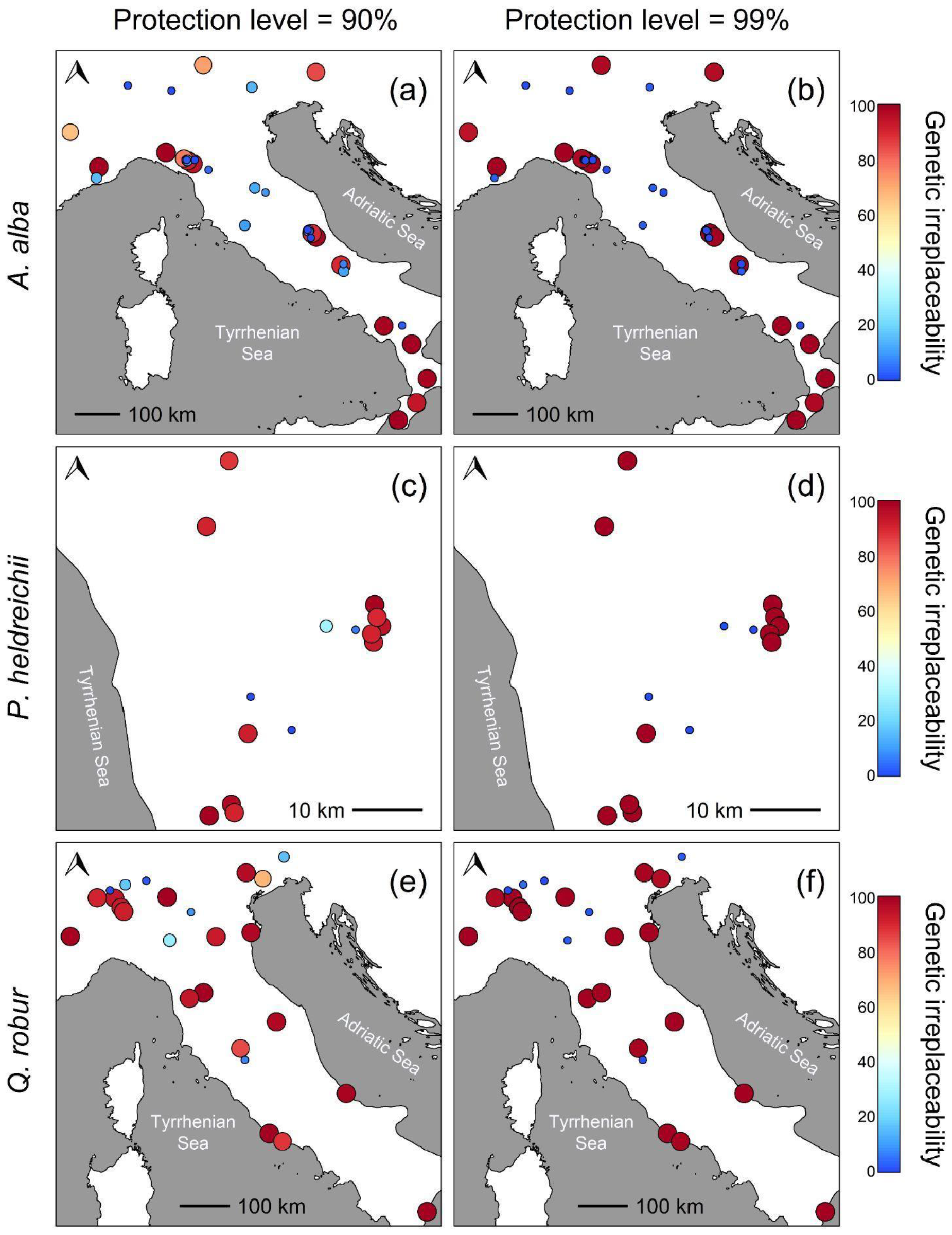
Genetic irreplaceability (GI) of sampled sites in *A. alba* (**a**,**b**), *P. heldreichii* (**c**,**d**), and *Q. robur* (**e**,**f**), as calculated over 100 replicates for different levels of IGD secured (i.e., 90% and 99% on the left and right columns, respectively). The GI of each site is indicated by both the size of the dots and their colour (a smaller size and blue tones indicate low GI, while a larger size and red tones indicate high GI).

### 3.2. Relationship between genetic irreplaceability and genetic diversity parameters

After multiple testing corrections, four out of the 15 tests used to assess the relationship between GI and genetic diversity parameters were statistically significant at a nominal α=0.05 (**Figure 4**). Particularly, *PAr* resulted to be associated with GI in all species, with the set of sites at higher GI showing a higher mean *PAr. PAr* displayed high variability in high GI sites (**Figure 4** and **Supplementary Table 2**). *A*_T_, *A*_S_ and *D*_A_ were not associated with GI. On the contrary, *Ne* turned out to be significantly associated with GI in *Q. robur*, where high GI sites had lower *Ne* on average compared to low GI sites.

**Figure 4.**
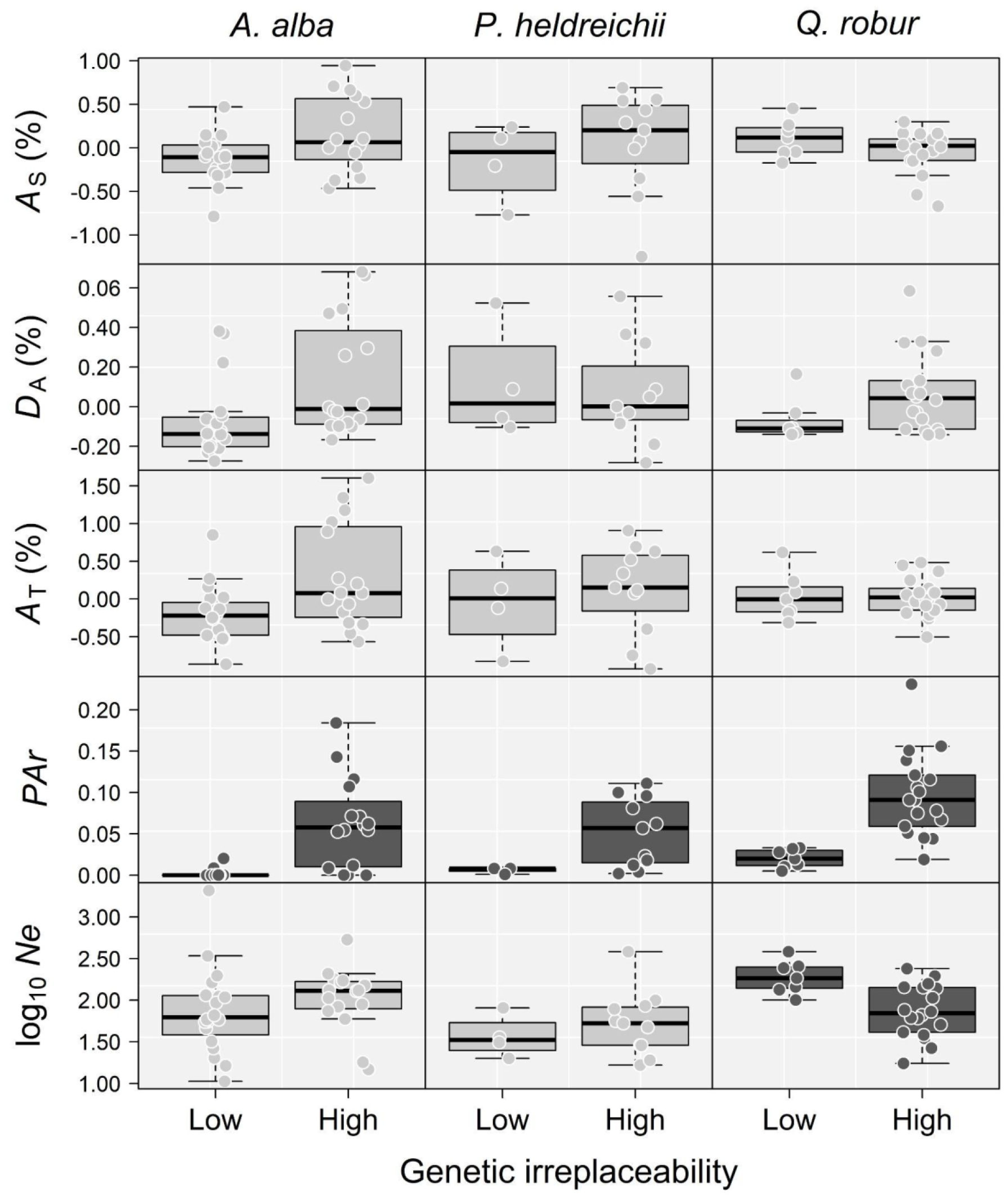
Distributions of site-specific contributions to within-site allelic diversity (*A*_S_), between-site allelic diversity (*D*_A_) and total allelic diversity (*A*_T_), private allelic richness (*PAr*) and effective population size (*Ne*) in sites with low and high genetic irreplaceability (GI<20% and GI>80%, respectively). Darker boxes indicate statistically significant differences between the means of the low and high GI sites after the Bonferroni correction for multiple testing. *Ne* values are presented on a logarithmic scale for graphical purposes.

## 4. Discussion

We applied SCP to intraspecific genetic data covering the Italian distribution of three forest tree species that are key elements of Mediterranean forest ecosystems (*Abies alba, Pinus heldreichii* and *Quercus robur*). Overall, we found that between 44% and 73% of the sampled sites, depending on the species, are needed to achieve adequate protection for all the alleles present in the sites studied. We showed that even if the objective was protecting fewer alleles (50% of alleles), the required number of sites did not change much. Moreover, even when the objective was to protect fewer alleles, the selected sites could ensure protection for a higher proportion of alleles. Finally, we found that private allelic richness (i.e., the number of alleles uniquely present in a given site) was the genetic parameter mostly related to genetic irreplaceability, a measure of the importance of conserving a given site to protect IGD. In the following, we discuss these results in the light of the conservation status of silver fir, Heldreich’s pine and pedunculate oak, as well as their life history traits and biogeographic history.

### 4.1. Full protection can be achieved at ‘little’ extra cost

Our SCP analysis resulted in a rather costly solution to protect IGD to the suggested level of 99% (Frankham, 2022). In fact, the percentage of protected sites was never less than 44% and included 16, 11 and 18 sites for silver fir, Heldreich’s pine and pedunculate oak, respectively. Nevertheless, a difference of only two to three sites separated the achievement of half and full protection of alleles, suggesting that low levels of protection (50% of alleles) are neither a parsimonious strategy for IGD conservation in these species, nor do they imply substantial cost reduction. Yet, any additional economic effort to achieve the 99% protection level (e.g., land acquisition and management) will depend critically on the available budget and socio-political will (Margules and Pressey, 2000; Naidoo et al., 2006).

### 4.2. Private allelic richness is not a perfect proxy for genetic irreplaceability

To explore the relationship between different approaches to priority setting, we related the genetic irreplaceability of a site to its relative contribution to within-site, between-site, and total allelic diversity, as well as to its effective population size and private allelic richness. These genetic parameters find wide application in conservation thanks to their sensitivity to demographic fluctuations (López-Cortegano et al., 2019; Petit et al., 1998; Tapio et al., 2006), and their capacity to comply with the principle of complementarity (Bonin et al., 2007) and to inform allocation of economic resources (Von Takach et al., 2023). The significant, positive correlation found between GI and *PAr* points to the spatial distribution of private alleles as one of the main genetic features driving site selection in our SCP analysis (**Supplementary Figure 1**). By relying on an SCP approach like the one used here, selection for sites enriched in private alleles has also been observed in other taxa, including freshwater fish species (Paz-Vinas et al., 2018) and marsupials (Von Takach et al., 2023).

However, such a correspondence between GI and *PAr* is not straightforward, as we found that some sites with low *PAr* can show a high GI. Thus, we recommend caution when using *PAr* as a perfect proxy for GI, as some highly irreplaceable sites could be ignored. We also note that selecting sites with private alleles implies including all non-private alleles contained in these sites in the final selection, which, depending on the spatial distribution of alleles and their frequencies, may increase the proportion of alleles that are adequately protected overall. Here, we never ended up in protecting less than 80% of the alleles, regardless of the level of protection set prior to the analysis.

### 4.3. Towards improving the interpretation and calculation of genetic irreplaceability

The other genetic parameters tested were not related to genetic irreplaceability, except for *Ne* in pedunculate oak, where high GI sites display a significantly lower *Ne* than low GI sites. This finding may suggest the use of genetic irreplaceability in combination with *Ne* to simultaneously locate priority sites for IGD conservation and evaluate their genetic health status, especially when dealing with highly fragmented populations as in the case of pedunculate oak in Italy (Avanzi et al., 2023). Rapidly shrinking populations in an area characterised by a strong genetic structure may explain the occurrence of high private allelic richness in sites with low *Ne* (see, for example, the case of the ‘Foglino’ and ‘Bosco Pantano’ sites in central and southern Italy, where *PAr* is high and *Ne* ranges from 27 to 81; **Supplementary Table 2**). On the other hand, new analyses with a broader species panel would be needed to clarify the relationship between *Ne* and GI and to provide guidance on how the genetic indicators adopted by the Convention on Biological Diversity (2022) can be harmonised with the SCP approach (in our case, for example, less than half of the high GI sites of each species had *Ne*>500, and our solutions often discarded sites with high *Ne*).

Our approach optimises the selection of sites so as to include at least 30% of the sites where an allele is present as an insurance against random allele loss. While valid, this approach ignores the potential relevance of within-site allele frequencies in calculating GI. Taking allele frequencies into account would be important from a conservation point of view because low frequency alleles are more prone to disappear due to the effects of genetic drift. For example, alleles with within-site frequency ≤5% were evenly distributed between low and high priority sites in the species under study (**Supplementary Figure 2**). The possible loss of rare alleles due to lack of protection could be overcome by setting specific targets for them, analogously to what is done when setting targets for species representation in conservation prioritisation (e.g., Rodrigues et al., 2004), or by ‘locking-in’ sites with higher proportions of rare alleles, so as to force their inclusion in the final selection of sites.

### 4.4. Glacial refugia are priority areas for conservation

The geographical range of the large majority of irreplaceable sites for *A. alba* corresponds to the Apennine mountain range, and includes all refugia areas that were proposed for this species based on both genetic and paleobotanical data (Magri et al., 2015; Piotti et al., 2017): the Calabrian arc (southern Apennines) including four sites; the central Apennines, including three sites located across two highly differentiated genetic lineages; and the northern Apennines, including four sites. Despite fragmentation and documented declines in abundance during the Holocene (Magri et al., 2015), such areas still maintain large population sizes, which will likely protect them from future genetic erosion (Forester et al., 2022; Kardos et al., 2021), and show high allelic diversity, local adaptation and growth vigour in provenance studies (Kerr et al., 2015; Piotti et al., 2017). The higher occurrence of high GI sites in the southern refugial area compared to the northern area may reflect high private allelic richness and a long history of genetic isolation from the northernmost populations (Piotti et al., 2017). Physical barriers (i.e., the Gran Sasso and Majella massifs) and possible constraints to long-distance dispersal due to heavy seeds and pollen (Leonarduzzi et al., 2016) may have acted in concert to prevent gene flow from the southern refugial area, facilitating the development of spatial genetic structuring (Piotti et al., 2017).

High GI sites for *Q. robur* also generally corresponded to refugial areas, which include the Pre-Alps, the southern slope of the Tuscan-Emilian Apennines, the Tyrrhenian coast and some areas in southern Italy, where the species was likely abundant at the onset of the Holocene (Magri et al., 2015). In particular, irreplaceable sites include six sites along the Po River (northern Italy), five sites in central Italy (two of them in Tuscany) and the putative refugium of ‘Bosco Pantano’ in the South. Contrary to what is observed in other refugial areas, ‘Bosco Pantano’ is genetically depauperate, with extremely low allelic diversity and effective population size (*Ne*<100) due to recent intensive human impact. The high irreplaceability of this site may rather reflect its extremely large genetic distinctiveness and the presence of private alleles (Avanzi et al., 2023). Similar to our findings, Beridze et al. (2023) have also recently shown that sites within refugial areas of *Castanea sativa* deserve high conservation priority.

### 4.5. Practical implications of SCP results

Based on comprehensive genetic datasets characterising neutral genetic diversity in a multi-refugial area, we were able to identify priority sites for conserving IGD in *A. alba, P. heldreichii* and *Q. robur*. The implications are manifold.

First, the genetic irreplaceability map (**Figure 3**) can make it possible to assess the current level of legal protection of sites whose loss would reduce the IGD of the species. All *P. heldreichii* sampled populations are within the Pollino National Park, whose perimeter was originally designed to include and protect all Italian populations of this species. The majority of *A. alba* and *Q. robur* populations are within National and Regional Parks, or public Reserves. Nonetheless, some irreplaceable sites for *Q. robur* are located on private property without any protection regime (e.g., the site with the highest genetic diversity in Italy; see Avanzi et al., 2023). Our results can contribute to change the protection status of such stands or to raise awareness among private owners toward a correct management of forest genetic resources. Finally, if some of these priority sites were ultimately unprotectable, replacements could be identified on the basis of their genetic irreplaceability, considering complementary genetic indicators (e.g., *Ne*), or through additional genetic sampling (Margules and Pressey, 2000).

Seed collection for *in situ* and *ex situ* conservation strategies as well as for reforestation programs, the latter having been massively funded in Italy in the last years (Mariotti et al., 2022), is usually carried out by the public sector in designated seed stands. Seed stands were not identified based on their genetic characteristics but mostly relying on the knowledge of local experts. Our results can help refine the Italian network of seed stands in terms of optimising the number and geographical position of selected stands (Wei and Jiang, 2020). For *A. alba*, for instance, >50 seed stands are registered at the national level (MASAF, 2023) but none is present in a relatively small genetic cluster found in central Italy (where two irreplaceable sites were identified) or in the site with the highest genetic diversity in Italy (‘Terranova di Pollino’ in Piotti et al., 2017), which we found to be irreplaceable starting from the lowest protection level. For *Q. robur*, there are about 100 seed stands registered, but none of them includes sites in central or southern Italy (the southernmost seed stand is at 43° 21’ latitude), where our SCP analysis identified six sites with high GI. This means that the genetic diversity of populations at the rear edge of the species’ distribution, which might have a disproportionate value in boosting the adaptive potential of the northernmost populations through gene flow or assisted migration (EUFORGEN, 2021), is currently not represented in seed collection initiatives in Italy. Finally, *P. heldreichii* has no registered seed stands so far, and our findings may be useful in defining an initial priority list.

The Pan-European network of genetic conservation units (Lefèvre et al., 2013; http://portal.eufgis.org/) shows a better agreement with our SCP analyses. For instance, the southernmost *Q. robur* population, which represents a unique genetic lineage and nowadays consists of only 64 individuals (Avanzi et al., 2023), is recognized as a genetic conservation unit but not as a seed stand. In addition, despite the absence of seed stands, a genetic conservation unit for *P. heldreichii* was established in a 166-ha area of the Pollino National Park that includes five of the irreplaceable sites identified in our analysis. Nonetheless, the selection of genetic conservation units (for which a first, partial genetic characterisation is currently underway in the framework of the EU-funded FORGENIUS project; www.forgenius.eu) could greatly benefit from SCP studies such as the one presented here.

### 4.6. Conclusions

SCP proved to be an effective method of identifying conservation areas capable of preserving the intraspecific allelic diversity of silver fir, Heldreich’s pine and pedunculate oak. In particular, we found that between 50% and 75% of sampled sites are required to protect IGD in these species, which is a higher percentage than the one usually recommended to protect habitats (30%; Convention on Biological Diversity, 2022). Importantly, we found that these percentages hold for both low (50%) and more ambitious percentages of alleles to be protected (>90%). Confirming theoretical expectations (Petit et al., 2003), glacial refugial areas were frequently included in the networks of proposed protected areas. This result may also apply to forest species for which SCP has not yet been undertaken.

Multifaceted applications of data and approaches presented here have been highlighted, with particular reference to optimising the seed collection necessary to fuel ongoing afforestation initiatives such as the commitment for planting at least 3 billion trees in the European Union by 2030 (EC, 2020). SCP could provide forest conservationists with clear guidance on which sites should be prioritised or added to the existing genetic conservation unit network. In this regard, informed decision-making is key to implementing any action aimed at achieving specific conservation targets (e.g., achieving 90-99% protection of existing forest genetic resources; Frankham, 2022; Hoban et al., 2023), monitoring IGD over time and flagging any need for active intervention (EUFORGEN, 2021).

Finally, we advocate that future research should focus on (i) describing the relative performance of SCP compared to more traditional methods based on allelic diversity and effective population size in different conservation scenarios, (ii) harmonising SCP objectives with the principles for the conservation of intraspecific genetic diversity proposed by the Convention on Biological Diversity (2022), and (iii) exploring the potential impact of sample size and the set of genetic markers used on SCP performance. In particular, the extension of SCP to genomic data should merit special attention to include both neutral and functional variants in the analysis (Hanson et al., 2020; Nielsen et al., 2023b, 2020; Xuereb et al., 2021), which promises to increase the effectiveness of genetic-informed conservation measures.

## Abbreviations

*A*_S_: the site-specific contribution to within-site allelic diversity
*D*_A_: the site-specific contribution to between-site allelic diversity
*A*_T_: the site-specific contribution to total allelic diversity
GI: genetic irreplaceability
IGD: intraspecific genetic diversity
*Ne*: effective population size
*PAr*: private allelic richness
SCP: spatial conservation planning

## Acknowledgments

E. Vajana is supported by the EU-funded project MedForAct (grant agreement ID 101107604). We thank all Parks and Reserves where sampling sites are located for permissions and logistic support. The acknowledgements to specific persons and institutions are reported in the articles where original data were published, indicated in the Materials & Methods section.

## Author contributions

**Elia Vajana**: Conceptualization; Data curation; Formal analysis; Visualization; Writing - original draft. **Marco Andrello**: Conceptualization; Formal analysis; Methodology; Supervision; Writing - review & editing. **Camilla Avanzi**: Conceptualization; Data curation; Formal analysis; Visualization; Writing - review & editing. **Francesca Bagnoli**: Funding acquisition; Writing - review & editing. **Giovanni G. Vendramin**: Funding acquisition; Supervision; Writing - review & editing. **Andrea Piotti**: Conceptualization; Data curation; Formal analysis; Funding acquisition; Supervision; Visualization; Writing - review & editing.

