## Supplementary Table 1 for "Spatial conservation planning of forest genetic resources in a Mediterranean multi-refugial area"

### Supplementary Materials

*Supplementary Table 1*

New *Abies alba* sites used in the present study other than those described in Piotti *et al*. (2017) and Santini *et al*. (2018). Sample size: number of individuals sampled.

| Site name | Site ID | Longitude | Latitude | Sample size |
|---|---|---|---|---|
| Gouta | GOU | 7.60633 | 43.94648 | 48 |
| Monte Orsaro | ORS | 9.9931 | 44.4151 | 48 |
| Monte Scala | SCL | 10.0406 | 44.3884 | 48 |
| Rocca Pianaccia | PIA | 10.0691 | 44.3811 | 48 |
| Lago Ballano | BAL | 10.0996 | 44.3631 | 48 |
| Monte Ventasso | VEN | 10.29177 | 44.38144 | 61 |
| Colle Romicito | CLR | 13.3972 | 42.6846 | 48 |

*Supplementary Table 2*

Site-specific contributions to within-site allelic diversity ($A_S$), between-site allelic diversity ($D_A$) and total allelic diversity ($A_T$) are reported together with private allelic richness (*PAr*), the point estimates of effective population size (*Ne*), and the lower and upper bounds of the 95% confidence intervals around the *Ne* estimates ($Ne_L$ and $Ne_H$, respectively). The genetic irreplaceability (GI) of each site is also shown for the 99% protection scenario, together with the site label, longitude ('Long.') and latitude ('Lat.').

| Site | Long. | Lat. | $A_S$ | $D_A$ | $A_T$ | *PAr* | *Ne* | $Ne_L$ | $Ne_H$ | GI |
|---|---|---|---|---|---|---|---|---|---|---|
| *Abies alba* | | | | | | | | | | |
| GOU | 7.60633 | 43.94648 | 0.0483 | -0.1702 | -0.1219 | 0.000206 | 397.2 | 111.6 | ∞ | 2 |
| PES | 7.66961 | 44.21128 | 0.0017 | -0.0027 | -0.001 | 0.008538 | 330.2 | 89 | ∞ | 100 |
| SAL | 6.89004 | 45.04732 | -0.3466 | 0.0121 | -0.3346 | 0 | ∞ | 150.4 | ∞ | 95 |
| TOC | 8.45981 | 46.16879 | -0.2347 | -0.2293 | -0.4641 | 0 | 1039.7 | 115.2 | ∞ | 0 |
| VBR | 9.65555 | 46.04139 | -0.2976 | -0.2325 | -0.5301 | 0.001269 | ∞ | 2085.4 | ∞ | 0 |
| SMO | 10.52305 | 46.65916 | -0.0055 | -0.1672 | -0.1727 | 0.000025 | ∞ | 179.7 | ∞ | 98 |

| | | | | | | | | | | |
|---|---|---|---|---|---|---|---|---|---|---|
| NOA | 11.84661 | 46.12929 | 0.0149 | -0.2095 | -0.1946 | 0 | ∞ | 340.3 | ∞ | 2 |
| TAR | 13.60088 | 46.48834 | 0.099 | -0.0206 | 0.0785 | 0.011475 | 166.6 | 83.7 | 1214.4 | 97 |
| NER | 9.50618 | 44.55988 | 0.3374 | -0.0622 | 0.2751 | 0.116281 | 474.4 | 131.6 | ∞ | 100 |
| ORS | 9.9931 | 44.4151 | -0.4646 | -0.101 | -0.5656 | 0.000156 | 19.5 | 14.7 | 26.3 | 98 |
| SCL | 10.0406 | 44.3884 | -0.2818 | -0.1941 | -0.4758 | 0 | 26.8 | 20.2 | 36.6 | 0 |
| RPM | 10.06333 | 44.38472 | -0.1329 | -0.2742 | -0.4071 | 0 | 72.7 | 46.1 | 141.6 | 2 |
| PIA | 10.0691 | 44.3811 | -0.283 | -0.1981 | -0.4811 | 0 | 37.1 | 26.4 | 55.6 | 0 |
| LAG | 10.0864 | 44.3689 | 0.0638 | -0.1631 | -0.0993 | 0.00135 | 406 | 111 | ∞ | 0 |
| BAL | 10.0996 | 44.3631 | -0.3737 | -0.0814 | -0.4551 | 0.054688 | 24.2 | 18 | 33.6 | 100 |
| VEN | 10.29177 | 44.38144 | -0.7879 | -0.0759 | -0.8638 | 0 | 14.1 | 10.6 | 18.7 | 0 |
| CER | 10.24157 | 44.28868 | 0.0287 | -0.0955 | -0.0668 | 0.061019 | 4251.2 | 164.2 | ∞ | 100 |
| ABE | 10.6671 | 44.14196 | 0.1455 | -0.1245 | 0.021 | 3.13E-05 | 94 | 55.5 | 230.9 | 0 |
| LVE | 11.93027 | 43.70861 | -0.3177 | -0.2054 | -0.5231 | 0.000181 | 111.1 | 58.4 | 452.4 | 2 |
| BTR | 12.22474 | 43.60015 | -0.4614 | -0.0616 | -0.523 | 0.000219 | 107.2 | 55 | 496.7 | 0 |
| PIG | 11.65699 | 42.81253 | 0.1485 | -0.1401 | 0.0085 | 0 | 3988.8 | 164.1 | ∞ | 2 |
| VDC | 13.37352 | 42.7053 | -0.1841 | -0.0829 | -0.2669 | 0 | 46.2 | 32 | 73.3 | 0 |
| CLR | 13.3972 | 42.6846 | -0.0913 | -0.0444 | -0.1357 | 0 | 104.3 | 58.7 | 309.9 | 0 |
| CEP | 13.43592 | 42.66952 | -0.0957 | -0.025 | -0.1207 | 0.0084 | 252.9 | 93.3 | ∞ | 2 |
| COR | 13.48965 | 42.62068 | -0.2183 | -0.0974 | -0.3157 | 0.0525 | 156.2 | 74 | 2734.9 | 100 |
| TOS | 13.60829 | 42.53076 | 0.1043 | -0.025 | 0.0793 | 0.107138 | ∞ | 208.2 | ∞ | 100 |
| SEG | 13.46444 | 42.50916 | -0.1105 | -0.1353 | -0.2458 | 0 | 21 | 16.3 | 27.5 | 0 |
| ABS | 14.28711 | 41.85733 | -0.0552 | 0.2591 | 0.2039 | 0.054688 | 1311.1 | 128.5 | ∞ | 100 |
| RSL | 14.35333 | 41.88305 | -0.0603 | 0.2225 | 0.1622 | 0.000131 | ∞ | 197.5 | ∞ | 0 |
| CME | 14.35638 | 41.70972 | -0.1024 | 0.3695 | 0.2671 | 0.000188 | 542.6 | 109 | ∞ | 2 |
| CIL | 15.45485 | 40.39774 | 0.5293 | 0.4936 | 1.0229 | 0.070813 | 92.1 | 59.6 | 176.8 | 100 |
| LAU | 15.95739 | 40.40635 | 0.4701 | 0.3816 | 0.8516 | 0.020119 | 110.2 | 66 | 265.3 | 0 |
| TDP | 16.21947 | 39.96029 | 0.9433 | 0.6613 | 1.6046 | 0.18425 | ∞ | 535.2 | ∞ | 100 |
| SIL | 16.63865 | 39.13343 | 0.7068 | 0.4714 | 1.1782 | 0.071306 | 222.2 | 105.9 | 8269.2 | 100 |
| SSB | 16.34803 | 38.55758 | 0.5985 | 0.2959 | 0.8944 | 0.062325 | 946.1 | 171.8 | ∞ | 98 |
| GAM | 15.85082 | 38.14041 | 0.665 | 0.6804 | 1.3454 | 0.143069 | 332.7 | 131 | ∞ | 100 |
| *Pinus heldreichii* | | | | | | | | | | |
| MAP | 15.96702 | 40.12763 | -1.2486 | 0.3213 | -0.9273 | 0.081 | 31.2 | 16.6 | 88 | 100 |

| | | | | | | | | | | |
|---|---|---|---|---|---|---|---|---|---|---|
| MLS | 15.92891 | 40.03959 | 0.0773 | 0.5564 | 0.6337 | 0.057 | 57.6 | 28.7 | 272.8 | 100 |
| SCR | 16.21158 | 39.93394 | -0.0092 | 0.0873 | 0.0781 | 0.111 | ∞ | 382.8 | ∞ | 100 |
| SCN | 16.21561 | 39.91712 | -0.3505 | -0.0478 | -0.3982 | 0.002 | 171.4 | 56.4 | ∞ | 100 |
| SCS | 16.22334 | 39.90519 | 0.2894 | 0.0492 | 0.3386 | 0.023 | 32.7 | 19 | 76.1 | 100 |
| SDO | 16.20669 | 39.89492 | 0.69 | 0.0017 | 0.6918 | 0.062 | ∞ | 99.7 | ∞ | 100 |
| MPO | 16.17967 | 39.90022 | -0.2061 | 0.0883 | -0.1178 | 0.008 | ∞ | 80.8 | ∞ | 0 |
| CPI | 16.21011 | 39.8834 | 0.5542 | -0.031 | 0.5232 | 0.004 | 208.9 | 78.8 | ∞ | 100 |
| TCA | 16.13025 | 39.90514 | 0.1106 | 0.5226 | 0.6333 | 0.008 | 94.1 | 35.7 | ∞ | 0 |
| CMC | 16.00335 | 39.81012 | 0.2399 | -0.104 | 0.1359 | 0.008 | 84.5 | 31.1 | ∞ | 0 |
| CDO | 15.99877 | 39.76088 | -0.5571 | -0.1896 | -0.7468 | 0.096 | 239.9 | 52.8 | ∞ | 100 |
| TPI | 16.07221 | 39.76538 | -0.77 | -0.0555 | -0.8255 | 0.001 | 34.1 | 20 | 72.7 | 0 |
| PIP | 15.97574 | 39.65416 | 0.4342 | -0.2829 | 0.1513 | 0.012 | 106.2 | 47.5 | ∞ | 100 |
| FAG | 15.93376 | 39.64975 | 0.5425 | 0.3651 | 0.9077 | 0.1 | ∞ | 86.1 | ∞ | 100 |
| ROS | 15.97077 | 39.66536 | 0.2033 | -0.085 | 0.1184 | 0.018 | 54.2 | 28.7 | 180.1 | 100 |
| *Quercus robur* | | | | | | | | | | |
| MER | 7.71317 | 44.78999 | -0.1453 | -0.115 | -0.2603 | 0.106 | 114.5 | 66.4 | 339.2 | 100 |
| LDS | 8.39053 | 45.44156 | 0.0848 | -0.0294 | 0.0554 | 0.051 | 655.4 | 142.6 | ∞ | 98 |
| TUR | 8.71116 | 45.5648 | 0.1183 | -0.121 | -0.0027 | 0.01 | ∞ | 244.1 | ∞ | 0 |
| FGN | 8.82743 | 45.43451 | 0.0616 | -0.1362 | -0.0747 | 0.059 | ∞ | 195.5 | ∞ | 100 |
| GER | 8.98614 | 45.27786 | 0.0876 | -0.1197 | -0.0321 | 0.044 | 97 | 60.8 | 214 | 100 |
| SIR | 9.05683 | 45.2102 | 0.1655 | -0.025 | 0.1404 | 0.067 | 1329.9 | 139.1 | ∞ | 100 |
| GRO | 9.09376 | 45.6609 | -0.0478 | -0.1059 | -0.1537 | 0.013 | ∞ | 382.4 | ∞ | 4 |
| PAL | 9.62455 | 45.7285 | 0.2015 | -0.1093 | 0.0922 | 0.005 | 521.5 | 133.8 | ∞ | 0 |
| CDC | 10.1484 | 45.45162 | -0.1494 | -0.061 | -0.2104 | 0.121 | 106.2 | 61.4 | 315.2 | 100 |
| FON | 10.74915 | 45.19796 | -0.0527 | -0.1332 | -0.186 | 0.02 | 272.1 | 100.5 | ∞ | 1 |
| MOR | 12.13213 | 45.86176 | -0.032 | 0.1085 | 0.0765 | 0.091 | 46 | 34.9 | 65 | 99 |
| CAV | 12.55146 | 45.76378 | 0.101 | -0.1138 | -0.0129 | 0.019 | ∞ | 239.6 | ∞ | 97 |
| FAG | 13.08088 | 46.13015 | -0.1723 | -0.1403 | -0.3126 | 0.028 | 1061.2 | 144.2 | ∞ | 2 |
| MES | 12.25539 | 44.85905 | -0.0062 | -0.1416 | -0.1478 | 0.091 | 3248.1 | 158.1 | ∞ | 100 |
| PFL | 11.37699 | 44.78402 | 0.0343 | 0.0512 | 0.0856 | 0.045 | 284.5 | 106.7 | ∞ | 99 |
| BDC | 10.20724 | 44.72643 | 0.2628 | -0.0313 | 0.2316 | 0.033 | ∞ | 256.2 | ∞ | 2 |
| MON | 10.70773 | 43.74992 | 0.0143 | 0.069 | 0.0833 | 0.078 | 163.4 | 76.1 | ∞ | 100 |

| | | | | | | | | | | |
|---|---|---|---|---|---|---|---|---|---|---|
| TAV | 11.06098 | 43.8419 | 0.1669 | 0.2813 | 0.4482 | 0.231 | 1348.2 | 143.7 | ∞ | 100 |
| FBN | 12.89251 | 43.35388 | 0.2997 | 0.0678 | 0.3675 | 0.139 | 54.8 | 38.1 | 91.8 | 100 |
| FAB | 11.98985 | 42.90642 | -0.5384 | 0.0345 | -0.5039 | 0.101 | 22.1 | 17.4 | 29 | 99 |
| ORV | 12.09845 | 42.71113 | 0.4537 | 0.1649 | 0.6187 | 0.032 | ∞ | 183.9 | ∞ | 0 |
| FOG | 12.71482 | 41.47387 | -0.08 | 0.3292 | 0.2492 | 0.156 | 36.4 | 26.6 | 54.2 | 100 |
| CRC | 13.04465 | 41.34019 | 0.1606 | 0.3227 | 0.4833 | 0.075 | 140.6 | 73 | 965.3 | 99 |
| POL | 14.6442 | 42.14798 | -0.318 | 0.1315 | -0.1864 | 0.116 | 59.8 | 41.1 | 101.5 | 100 |
| PAN | 16.68015 | 40.15543 | -0.6705 | 0.5848 | -0.0857 | 0.151 | 63.2 | 50.9 | 81.2 | 100 |

*Supplementary Figure 1*

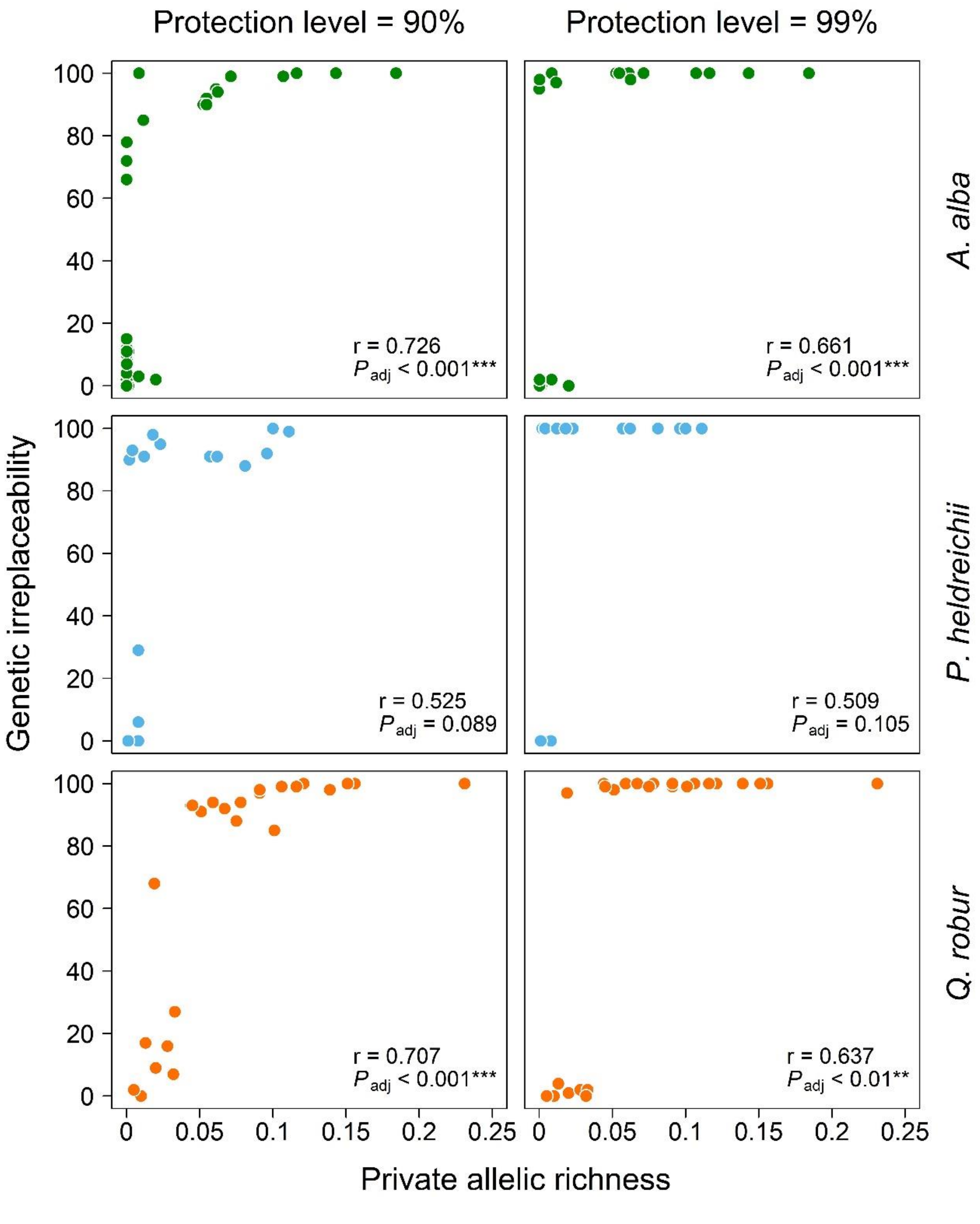


Relationship between private allelic richness (*PAr*; *X*-axis) and genetic irreplaceability (GI; *Y*-axis) under the 90% protection level (left column) and the 99% protection level (right column). Results for *A. alba* are reported in the first row, for *P. heldreichii* in the second row, and for *Q. robur* in the third row. For all species and both protection levels, GI increases with increasing *PAr*. Statistically significant, positive correlations were found for *A. alba* and *Q. robur* (r: Pearson's correlation coefficient). A non-linear relationship between the variables can be observed. To reduce the inflation of the familywise type I error, the Bonferroni correction was applied to the *p*-values of the correlation tests involving the same species. A test was considered statistically significant if its adjusted *p*-value was equal to or less than the nominal significance threshold of 0.05.

*Supplementary Figure 2*

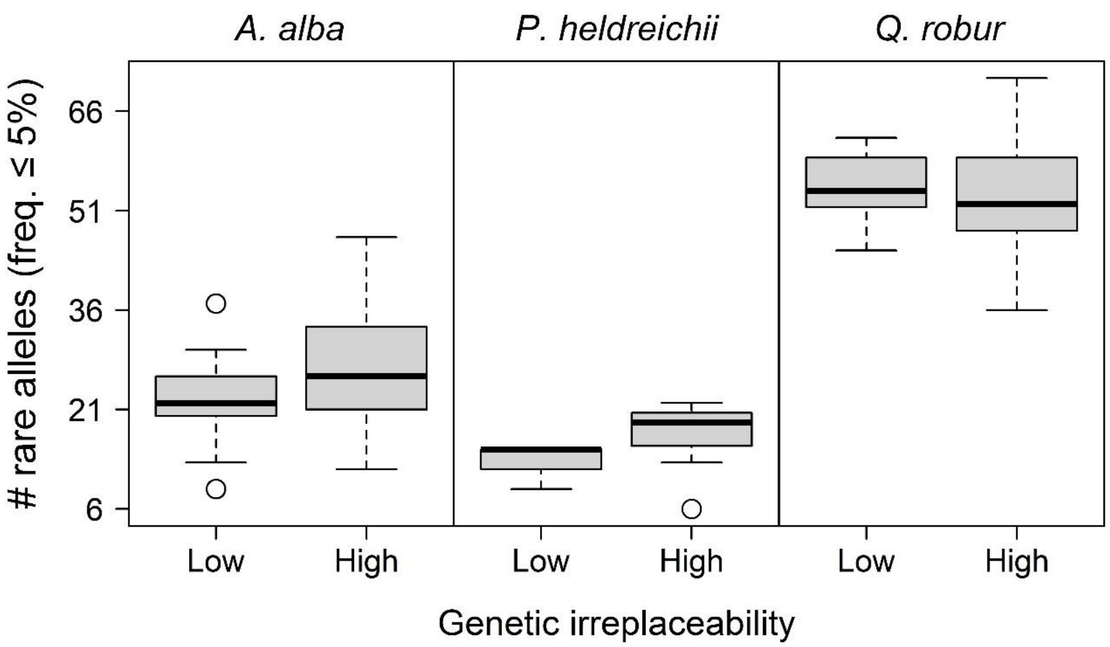


Distribution of rare alleles (defined here as alleles with within-site frequency ≤5%) in the low and high genetic irreplaceability sites (GI<20% and GI>80%, respectively). To test whether there was a significant difference in the amount of rare alleles occurring in the low versus high priority sites, a series of *t*-tests (one per species) were performed after testing for homogeneity of variances using a Levene's test. No significant difference was observed between the means of low and high priority sites at a nominal significance threshold of 0.05, indicating that the SCP procedure used here does not prioritise sites enriched in rare alleles. This relationship was assessed using the GI values obtained when protecting 99% of the alleles.
